# ARF1 dimerization is essential for vesicle trafficking and dependent on activation by ARF-GEF dimers in Arabidopsis

**DOI:** 10.1101/2020.01.21.913905

**Authors:** Sabine Brumm, Mads Eggert Nielsen, Sandra Richter, Hauke Beckmann, York-Dieter Stierhof, Manoj K. Singh, Angela-Melanie Fischer, Venkatesan Sundaresan, Gerd Jürgens

**Author notes:** Corresponding author: Gerd Jürgens, Center for Plant Molecular Biology (ZMBP), Developmental Genetics, University of Tübingen, Auf der Morgenstelle 32, 72076 Tübingen, Germany, ORCID: 0000-0003-4666-8308. **Material distribution footnote** The author responsible for distribution of materials integral to the findings presented in this article is: Gerd Jürgens.

## Abstract

Membrane traffic maintains the organization of the eukaryotic cell and delivers cargo proteins to their subcellular destinations such as sites of action or degradation. Membrane vesicle formation requires ARF GTPase activation by the SEC7 domain of ARF guanine-nucleotide exchange factors (ARF-GEFs), resulting in the recruitment of coat proteins by GTP-bound ARFs. *In vitro* exchange assays were done with monomeric proteins, although ARF-GEFs have been shown to form dimers *in vivo*. This feature is conserved across the eukaryotes, however its biological significance is unknown. Here we demonstrate ARF1 dimerization *in vivo* and we show that ARF-GEF dimers mediate ARF1 dimer formation. Mutational disruption of ARF1 dimers interfered with ARF1-dependent trafficking but not coat protein recruitment in Arabidopsis. Mutations disrupting simultaneous binding of two ARF1•GDPs by the two SEC7 domains of GNOM ARF-GEF dimer prevented stable interaction of ARF1 with ARF-GEF and thus, efficient ARF1 activation. Our results suggest a model of activation-dependent dimerization of membrane-inserted ARF1•GTP molecules required for coated membrane vesicle formation. Considering the evolutionary conservation of ARFs and ARF-GEFs, this initial regulatory step of membrane trafficking might well occur in eukaryotes in general.

## Introduction

Activation of small GTPase ARF1 by guanine-nucleotide exchange factors (ARF-GEFs) plays a pivotal role in membrane traffic across the eukaryotes (Donaldson and Jackson, 2011). GDP-bound ARF1 interacts with the catalytic SEC7 domain of ARF-GEFs on donor membranes, resulting in GDP-GTP exchange on ARF1 and membrane insertion of its myristoylated N-terminal hasp (Casanova, 2007; Anders et al., 2008a; Bui et al., 2009). GTP-bound ARF1 interacts with coat proteins involved in vesicle formation and cargo recruitment (D’Souza-Schorey and Chavrier, 2006; Gillingham and Munro, 2007; Singh and Jürgens, 2018). ARF1•GTP forms dimers *in vitro*, which are required for scission of membrane vesicles from donor membrane (Beck et al., 2008 and 2011). ARF1 dimer formation is disrupted by a Y_35_A mutation, which reduces the yield of vesicles dramatically *in vitro* and fails to complement the lethality of *arf1 arf2* mutant yeast (Beck et al., 2008). How ARF1 dimers form *in vivo* has not been addressed but might be related to ARF-GEF action. Large ARF-GEFs such as human GBF1 or Arabidopsis GNOM have a stereotypic domain organization including an N-terminal dimerization (DCB) domain (Casanova, 2007; Anders et al., 2008; Bui et al., 2009). The DCB domain can interact with another DCB domain and with at least one other ARF-GEF domain (Grebe et al., 2000; Ramaen et al., 2007; Anders et al., 2008b). Although conserved across the eukaryotes, the biological significance of ARF-GEF dimerization is not known. Our results presented here suggest that ARF-GEF dimers generate ARF1•GTP dimers during the activation process, allowing productive vesicle formation.

## Results and Discussion

### In-vivo occurrence and biological significance of ARF1 GTPase dimers

To address the issue of ARF1 dimerization, we tested ARF1 wild-type and two variants – activation-deficient ARF1-T_31_N and hydrolysis-deficient ARF1-Q_71_L (Dascher and Balch, 1994; Singh and Richter et al., 2018) – for interaction by co-immunoprecipitation (for overview of mutant proteins see Supplemental Figure 1). Both ARF1-T_31_N and ARF1-Q_71_L co-precipitated endogenous ARF1, although ARF1 wild-type failed to do so (Figure 1A). However, ARF1-T_31_N strongly interacted with ARF-GEF GNOM whereas ARF1-Q_71_L did not, suggesting that ARF1•GDPs might be bridged by ARF-GEF dimer whereas ARF1•GTPs might display interaction independently of ARF-GEF. To test this idea, we made use of the putatively dimerization-deficient ARF1-Y_35_A mutant (Beck et al., 2008). We generated transgenic Arabidopsis lines that inducibly co-expressed either ARF1-Y_35_A,Q_71_L-GFP and ARF1-Y_35_A,Q_71_L-RFP or ARF1-Q_71_L-GFP and ARF1-Q_71_L-RFP. FRET-FLIM measurements revealed interaction between ARF1-Q_71_L proteins in seedling root cells, which was prevented by the additional Y_35_A mutation (Figure 1B). These data suggest that membrane-bound ARF1•GTP molecules form dimers by direct physical interaction.

**Figure 1.**
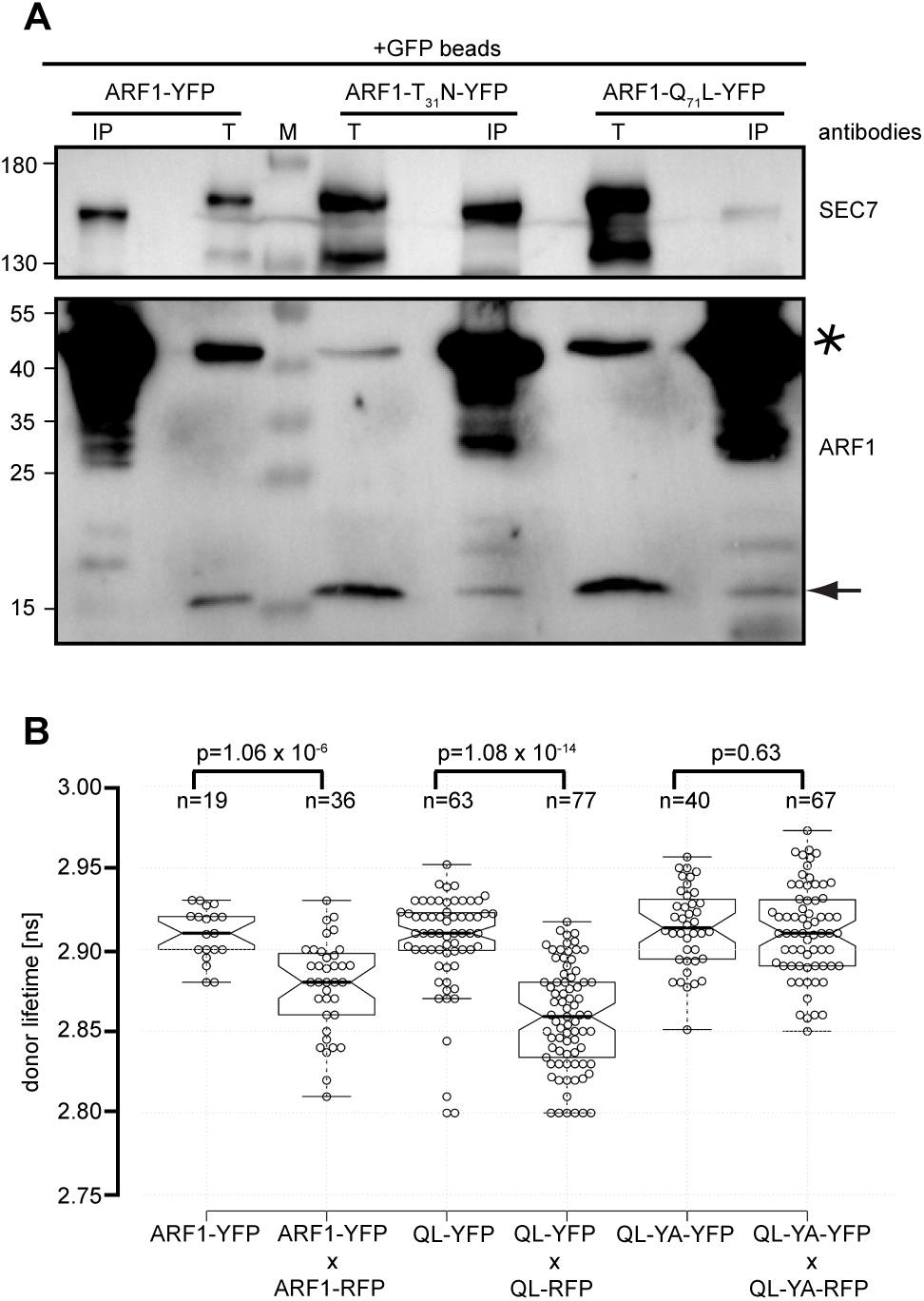
In-vivo interaction between ARF1•GTP molecules revealed by co-immunoprecipitation and FRET-FLIM analysis. **(A)** Estradiol-inducible (20µM 7h) expression of ARF1-YFP, activation-deficient ARF1-T_31_N-YFP and hydrolysis-deficient ARF1-Q_71_L-YFP, immunoprecipitation with anti-GFP beads from transgenic Arabidopsis seedling extracts, and immunoblotting of PAGE-separated precipitates with anti-SEC7(GNOM) and anti-ARF1 antisera. Antisera indicated on the right. T, total extract; IP, immunoprecipitate; M, molecular markers (sizes in kDa indicated on the left). Asterisk, ARF1-YFP fusion proteins; arrow, endogenous ARF1 (both detected with anti-ARF1 antiserum); SEC7, antiserum detecting SEC7 domain of GNOM. **(B)** FRET-FLIM analysis of ARF1-ARF1 interaction in Arabidopsis seedling root cells after estradiol induction (20µM 4h). The life time of hydrolysis-deficient ARF1-Q_71_L-YFP (QL-YFP) was reduced whereas ARF1-Q_71_L-YFP bearing dimerization-disrupting Y_35_A mutation (QL-YA-GFP) showed normal FRET-FLIM ratios. For comparison, the life time of ARF1-YFP was slightly reduced. Box plots of donor life time in ns of at least 19 independent measurements for each sample (exact numbers are indicated by n). Medians are represented by the center lines and notches indicate 95% confidence interval. Tukey whiskers extend to the 1.5xIQR and data points are plotted as bee swarm. Exemplary p values (two tailed t-test assuming equal variances, alpha=0.05) are indicated in the graph.

To examine the biological significance of ARF1 dimers, we analyzed the ability of ARF1-Y_35_A to rescue the secretion of alpha-amylase from tobacco protoplasts inhibited by ARF1-T_31_N expression (Figure 2A and 2B).

**Figure 2.**
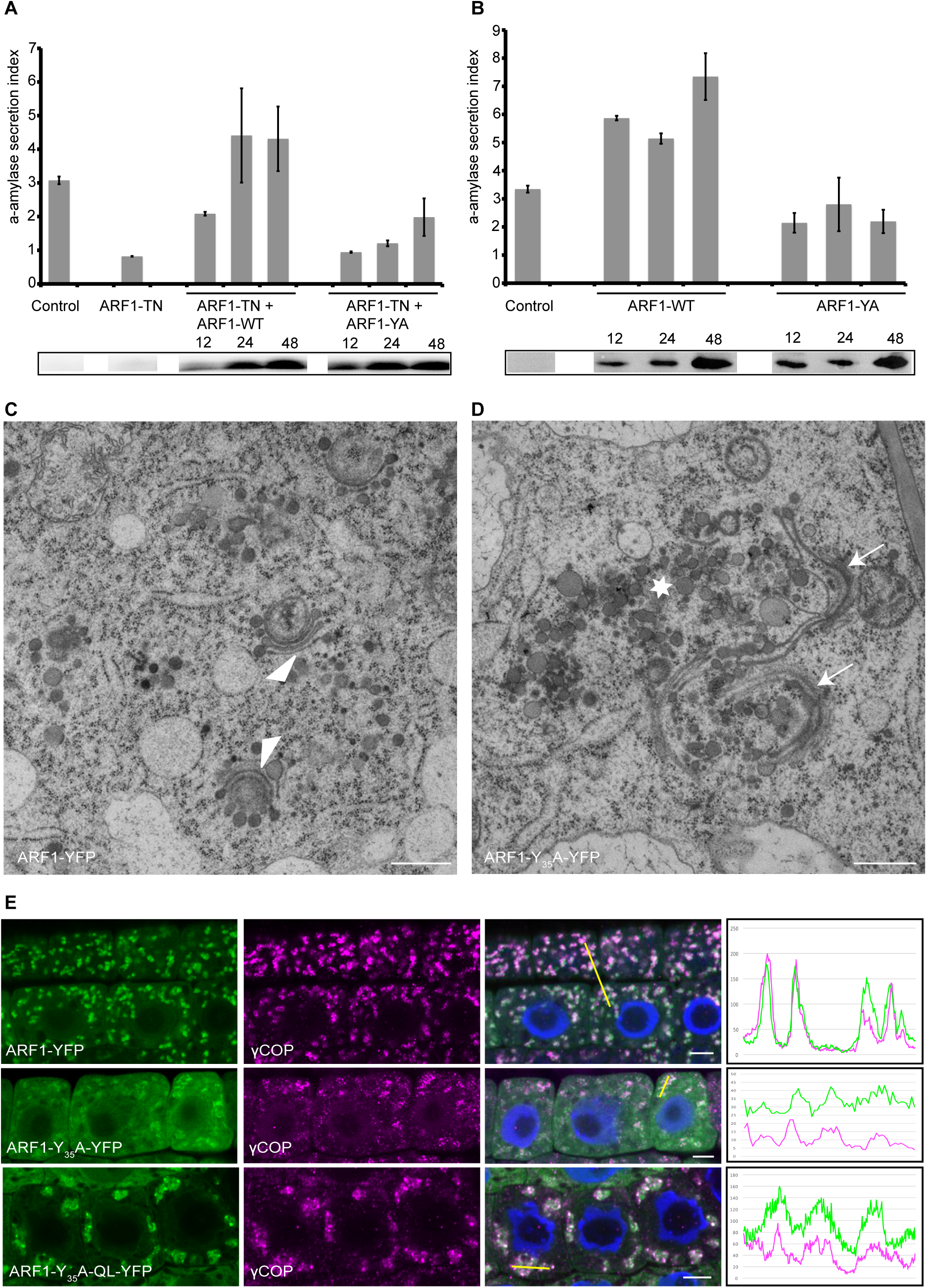
Biological consequences of dimerization-deficient ARF1 expression. (**A**, **B**) Secretion of alpha-amylase from tobacco protoplasts (A) inhibited by ARF1-T_31_N (TN) was restored by overexpression of ARF1 (WT) but not ARF1-Y_35_A (Y35A) and (B) impaired by ARF1-Y_35_A (Y35A) compared to wild-type control. Bottom panels: Antibody detection of GFP linked to ARF1 expression. (**C**, **D**) Electron microscopic analysis of epidermal cells at the upper end of the seedling root meristem expressing ARF1-YFP (C) or ARF1-Y_35_A-YFP (D) in response to 10 µM estradiol for 4 h. Golgi stacks (arrowheads) were bent in (C) but replaced by clusters of interconnected membrane vesicles (asterisk) and Golgi remnants (arrows) in (D). Scale bars, 500 nm. (**E**) Immunostaining of COPI subunit γCOP in seedling root cells expressing ARF1-YFP, ARF1-Y_35_A-YFP or ARF1-Y_35_A,Q_71_L-YFP in response to 20 µM estradiol for 4h. ARF1 variant (green), γCOP (magenta), merged images with DAPI-stained nuclei (blue). Co-localization of ARF1 and γCOP in regions of interest (ROI; yellow lines) shown in line intensity profiles. Scale bar, 5 µm.

Rising concentrations of co-expressed wild-type form of ARF1 overcame the inhibition by ARF1-T_31_N. In contrast, co-expression of comparable concentrations of ARF1-Y_35_A largely failed to restore alpha-amylase secretion (Figure 2A). In addition, strong expression of ARF1-Y_35_A interfered with alpha-amylase secretion on its own (Figure 2B). Thus, ARF1 dimerization is required for ARF1-dependent membrane trafficking. We also analyzed the consequences of ARF1-Y_35_A overexpression in seedling root cells at the ultrastructural level (Figure 2C and 2D). ARF1-Y_35_A disrupted Golgi organization, resulting in strings of interconnected membrane vesicles, whereas overexpression of ARF1 wild-type protein only caused slight bending of the Golgi stacks. However, overexpression of ARF1-Y_35_A did not interfere with membrane recruitment of COPI subunit γCOP (Figure 2E). Thus, ARF1 dimerization is essential for membrane trafficking.

### Regulation of ARF1 dimer formation

How could ARF1 dimer formation be regulated? One candidate is the activating ARF-GEF which itself forms dimers (Ramaen et al., 2007; Anders et al., 2008). ARF1•GDP binding by ARF-GEF involves the C-terminal loop after helix J (loop>J) of the SEC7 domain of human GBF1, as demonstrated by specific mutations that interfere with ARF1 binding (Lowery et al., 2011). We introduced homologous mutations into Arabidopsis GNOM to generate GN-loop>J(3A) mutant protein (Figure 3A). To assess the biological consequences of the *GN-loop>J(3A)* mutation, we analyzed the mutant phenotypes of plants expressing the GN-loop>J(3A) mutant protein in various *gnom* mutant backgrounds (Figure 3B-G).

**Figure 3.**
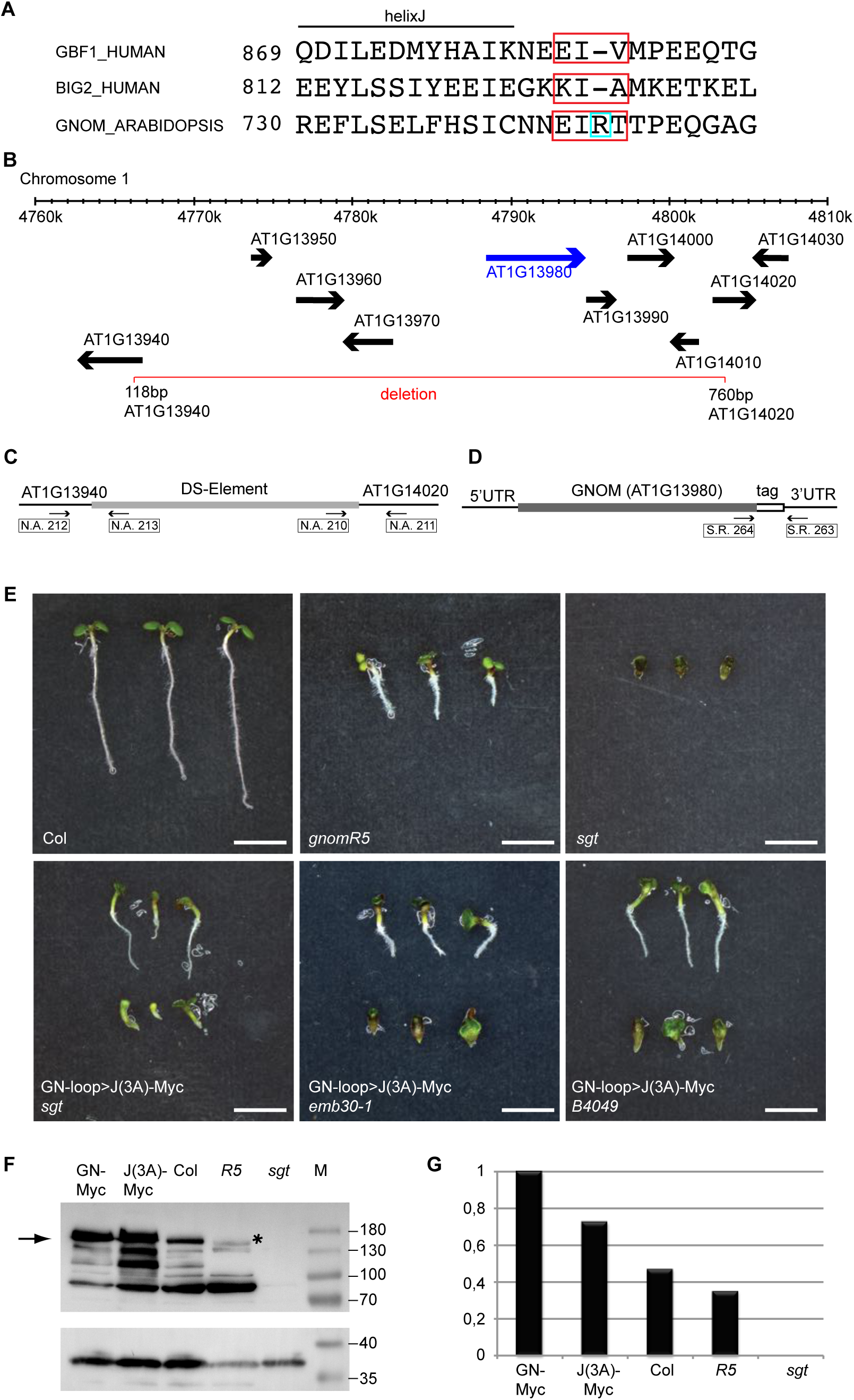
Rescue activity of GN-loop>J(3A)-Myc in *gnom-sgt* deletion and other *gnom* mutants. (A) Alanine substitution sites (red boxes) in the loop after helix J (loop>J) of SEC7 domain of ARF-GEFs human GBF1 (Lowery et al., 2011), human BIG2 (Lowery et al., 2011) and Arabidopsis GNOM. In GN-loop>J(3A) mutant protein, amino acid residues 744 to 747 (EIRT) are replaced by AARA. (B) Diagram of 50 kb genomic segment of chromosome 1 displaying GNOM and adjacent genes (arrows pointing towards 3’ end). The *GNOM* gene is highlighted in blue. The straddling 37 kb deletion (named *gnom-sgt*) encompassing GNOM and 8 flanking genes is indicated by a red line (Kumaran et al., 1999). End points of deletion are indicated by basepair (bp) numbers of respective genes. (**C, D**) Primer combinations for genotyping seedlings to detect (C) the *gnom-sgt* deletion or (D) the endogenous *GNOM* gene and a *GNOM* transgene encoding a C-terminally tagged protein. (**E**) Wild-type seedlings (Col), *gnom-sgt* deletion seedlings (*sgt*) and partially rescued *gnom-sgt* deletion seedlings bearing a *GN-loop>J(3A)-Myc* transgene; seedlings homozygous for the weak *gnom-R5* allele are shown for comparison. Partial rescue of exchange-deficient *gnom* seedlings (*emb30*) or membrane-assocation-deficient *gnom* seedlings (*B4049*). Scale bars, 2.5 mm. (**F, G**) GNOM protein expression levels of wild-type (Col), *gnom* mutant allele *R5*, *gnom-sgt* deletion (*sgt*), and *GNOM* transgenes *GNOM-myc* (GN-Myc) and *GN-loop>J(3A)* (J(3A)-Myc)) detected by anti-SEC7 domain antiserum. Loading control: unstripped membrane re-probed with anti-SYP132 antiserum. (F) Immunoblot; M, marker lane; protein sizes in kDa on the right. Arrow, GNOM band at 165 kDa; asterisk, truncated GNOM protein of *gnom-R5* at 155 kDa. (G) Normalized expression levels; GNOM from *GN-myc* set at 1. Col, wild-type level of GNOM protein. Note that *gnom-sgt* deletion zygotes complete embryogenesis and give rise to highly abnormal seedlings because the retrograde COPI traffic from Golgi to ER is jointly mediated by GNOM and the paralogous ARF-GEF GNL1 whereas the GNOM-mediated polar recycling of auxin efflux carrier PIN1 from endosomes to the basal plasma membrane cannot be mediated by GNL1 (Richter et al., 2007). Note also that the mutations *emb30* and *B4049* both reside in the SEC7 domain of GNOM. Allele *emb30* codes for a catalytically inactive E_658_K mutant protein, resulting in grossly abnormal seedlings (Meinke, 1985; Mayer et al. 1993; Shevell et al., 1994). *B4049* codes for a G_579_R mutant protein that is still catalytically active but fails to associate with endomembranes because the DCB-ΔDCB interaction is compromised, which also results in grossly abnormal seedlings (Anders et al., 2008b; Busch et al., 1996).

The mutant protein displayed some residual activity, very incompletely rescuing the *gnom-sgt* deletion which spans *GNOM* and 4 adjacent genes on either side (Figure 3B and 3E). Interestingly, GN-loop>J(3A) mutant protein was not able to completely rescue *gnom* mutant alleles *emb30* and *B4049*, whereas the latter two complemented each other (Figure 3E; Busch et al., 1996; Anders et al., 2008). The rescued seedlings rather resembled phenotypically seedlings bearing the weak allele *gnomR5* (Geldner et al., 2004). Whereas *gnomR5* encodes C-terminally truncated GNOM protein that accumulated to reduced level compared to wild-type, GN>loop>J(3A) accumulated to higher level than wild-type, suggesting a primary defect in protein function rather than protein or RNA stability (Figure 3F and 3G). A more detailed phenotypic analysis of incompletely rescued seedlings revealed *gnomR5*-like defects in cotyledon vasculature, lateral root initiation and primary root growth, regardless of the *gnom* mutant background (Figure 4A-C).

**Figure 4.**
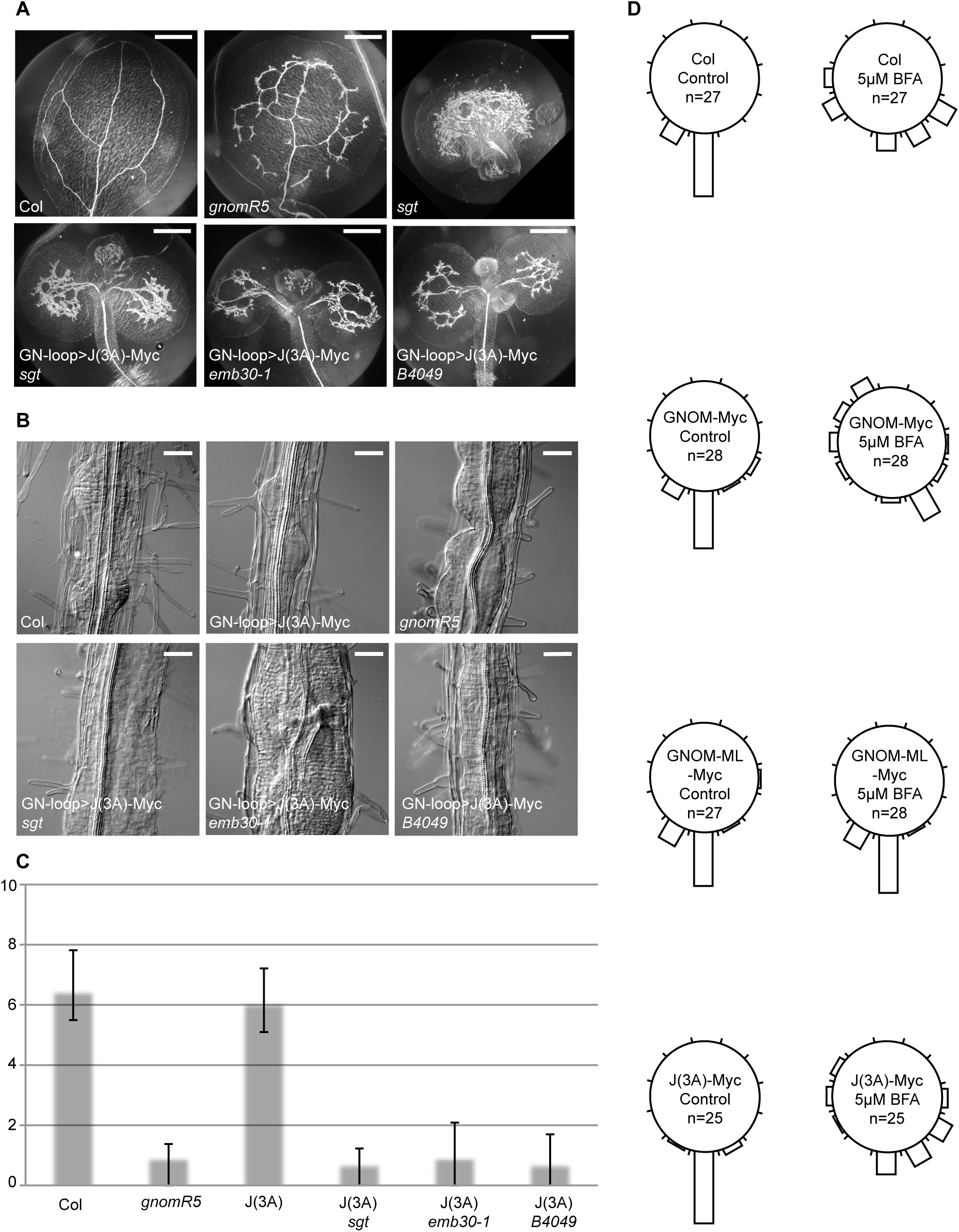
Seedling phenotypes of *gnom* mutants bearing the *GN-loop>J(3A)-Myc* transgene. (A) Vascular tissue differentiation in cotyledon. Scale bars, 500 µm. (B) Lateral root initiation. Scale bars, 50 µm. (C) Primary root length (in mm). J(3A), GN-loop>J(3A) in wild-type background or in different *gnom* mutant genotypes indicated (*sgt*, *emb30-1*, *B4049*). (D) Root gravitropism in seedlings treated with 5 µM BFA and untreated control seedlings. GN-loop>J(3A)-myc (J(3A)) in wild-type background behaves like Col or GN-myc. Col, wild-type; *gnomR5*, weak mutant allele; J(3A), GN-loop>J(3A) in wild-type background; *sgt*, *gnom-sgt* deletion; *emb30-1*, catalytically defective gnom-emb30; *B4049*, membrane-association-defective gnom-B4049; *GNOM-Myc*, *GNOM-Myc* transgene; *GNOM-ML-Myc*, engineered BFA-resistant GNOM; *J(3A)-Myc*, *GN-loop>J(3A)-Myc* transgene.

In the wild-type background, GN>loop>J(3A) had no noticeable phenotypic effects e.g. in primary root growth or root gravitropic response and displayed BFA-sensitivity, like wild-type (Figure 4C and D). Postembryonally, *GN-loop>J(3A)* plants had a twisted rosette, similar to *emb30/B4049* trans-heterozygous plants, but subsequently grew to normal height during flowering (Figure 5).

**Figure 5.**
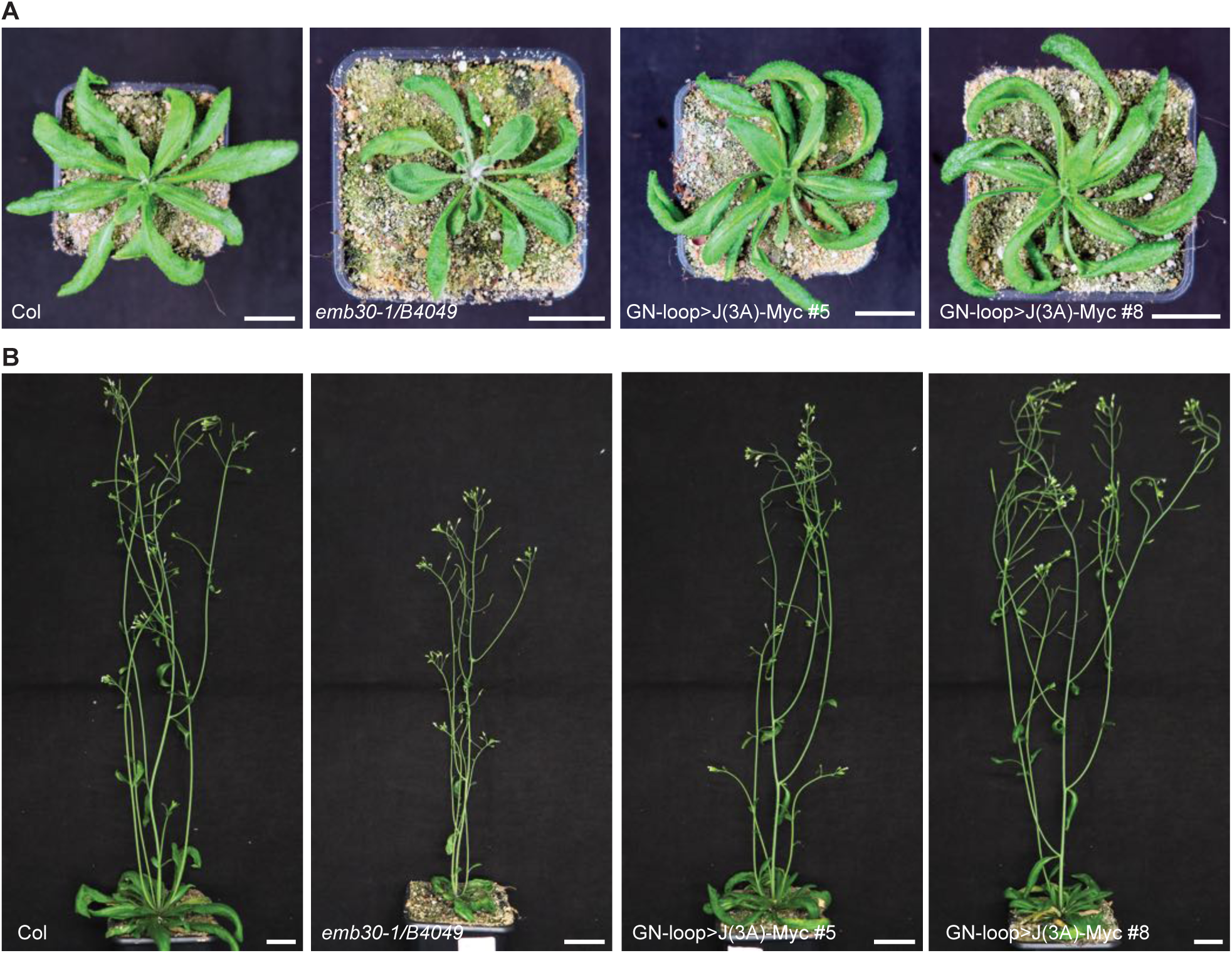
Developmental phenotypes of wild-type plants expressing GN-loop>J(3A)-Myc. (A) Rosette stage. Col, wild-type. Note slightly twisted rosettes of *trans*-heterozygous plants bearing nearly fully complementing *gnom* alleles (*emb30-1/B4049*) and of *GN-loop>J(3A)-Myc* transgene in Col-0 (two transgenic lines #5, #8). (B) Plants after the onset of flowering. Same genotypes as in (A). Note nearly normal stature of *GN-loop>J(3A)-Myc* transgenic plants. Scale bars, 2 cm.

In conclusion, the GN>loop>J(3A) mutant protein seems to retain residual activity but cannot be complemented by previously defined GNOM mutant proteins with specific defects in membrane association (B4049) or GDP-GTP exchange activity (emb30).

Because the human ARF-GEF GBF1 with a J-loop mutation displayed impaired ARF1 binding (Lowery et al., 2011), we addressed this issue, using co-immunoprecipitation assays. ARF1-YFP was co-immunoprecipitated with Myc-tagged GNOM wild-type protein but not with Myc-tagged GN-loop>J(3A) mutant protein, using anti-YFP or anti-Myc beads (Figure 6A and 6B). This compromised interaction between GN-loop>J(3A) and ARF1•GDP was consistent with a mutant phenotype corresponding to low level of ARF-GEF activity of GNOM as described above (see Figure 3B-G and Figure 4). The phenotypically detectable residual activity of GN-loop>J(3A) mutant protein suggested that its strongly reduced interaction with ARF1 appeared to be below the detection limit. We thus stabilized the presumed interaction of the mutant ARF-GEF with ARF1 by estradiol-induced expression of activation-deficient ARF1-T_31_N-YFP, because the T_31_N mutation blocks the GDP-GTP exchange (Dascher and Balch, 1994). This change revealed interaction between GN-loop>J(3A) protein and ARF1 in the co-immunoprecipitation assay (Figure 6C). This result suggested that ARF1 binding was impaired but the GN-loop>J(3A) protein was still able to carry out the GDP-GTP exchange, consistent with the partial rescue of the *gnom-sgt* deletion.

**Figure 6.**
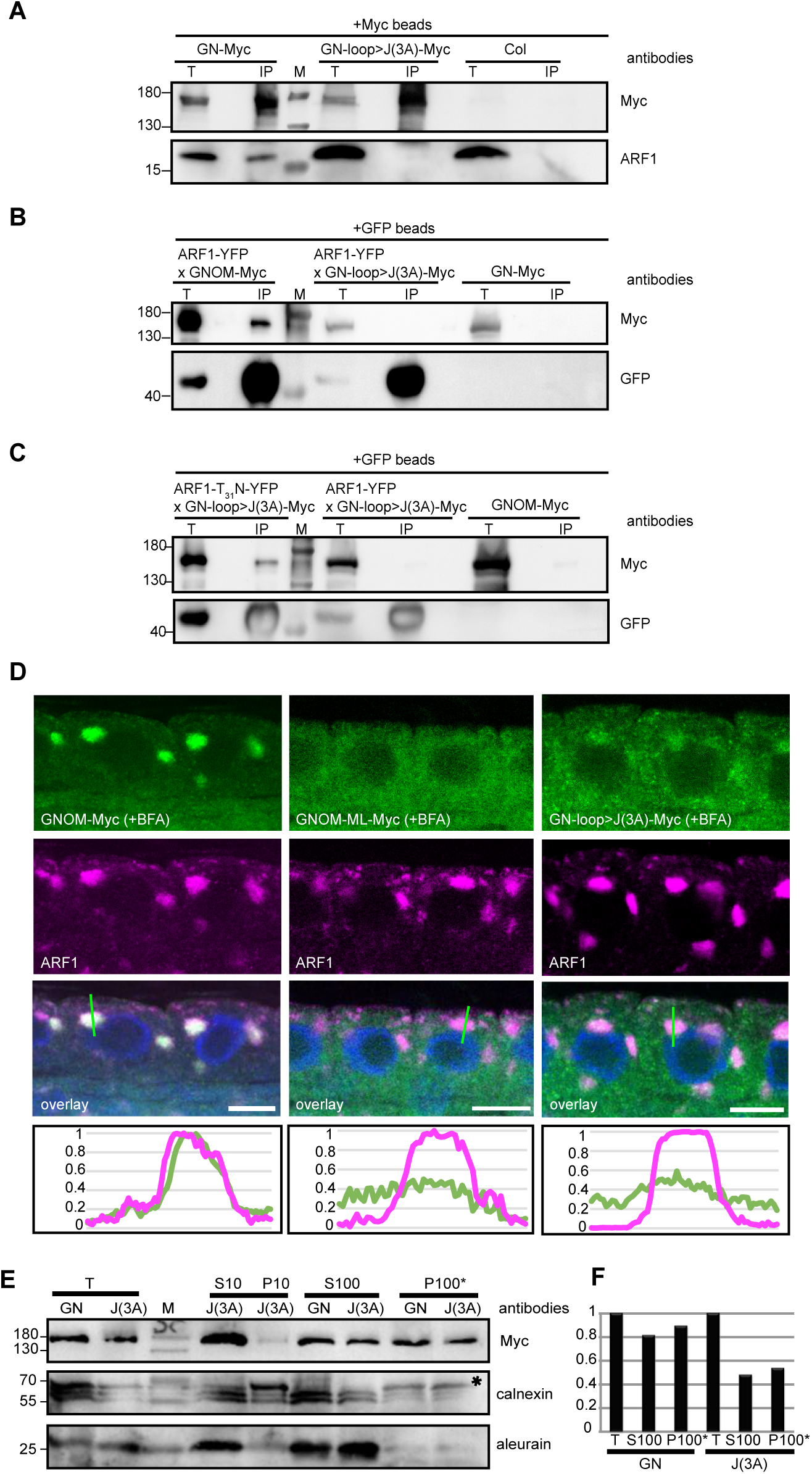
ARF-GEF GN-loop>J(3A) mutant protein: interaction with ARF1, subcellular localization and membrane association. (**A-C**) Co-immunoprecipitation from Arabidopsis seedling extracts. No detectable interaction of (A) endogenous ARF1 or (B) YFP-tagged ARF1 with GN-loop>J(3A)-Myc compared to GNOM-Myc wild-type control, following IP with (A) anti-Myc beads or (B) anti-GFP beads. (C) Activation-deficient ARF1-T_31_N-YFP yielded co-IP signal of GN-loop>J(3A)-Myc; IP with anti-GFP beads. Col, Columbia wild-type control. T, total extract; IP, immuneprecipitate; M, molecular markers (sizes in kDa, *left*). (**D**) Immunostainings of BFA-treated seedling roots. GNOM co-localized with ARF1, GN-loop>J(3A) essentially behaved like engineered BFA-resistant GNOM-ML, not accumulating on the ARF1-positive endomembrane. Line scans are indicated with green lines. (**E**, **F**) Cell fractionation revealed comparable partitioning between cytosol and membrane of Myc-tagged GNOM wild-type (GN) and Myc-tagged GN-loop>J(3A) mutant protein (J(3A)). (E) Immunoblot with antisera indicated on the right (controls: calnexin [asterisk], membrane protein (Huang et al., 1993); AALP, soluble protein (Holwerda et al., 1990); M, molecular markers (sizes in kDa, *left*). (F) Quantitation of anti-Myc signal intensities; total extracts set at 1. T, total extract; S10 and P10, supernatant and pellet of 10,000 *g* centrifugation; S100 and P100*, supernatant and washed pellet of 100,000 *g* centrifugation.

The fungal toxin brefeldin A (BFA) inhibits the exchange reaction, thus stabilizing abortive complexes of ARF-GEF and ARF1•GDP on endomembranes (Geldner et al., 2003; Mossessova et al., 2003; Renault et al., 2003). Treating seedling roots with BFA resulted in co-localization of GNOM and ARF1 in endosomal membrane aggregates called BFA compartments (Figure 6D) (Geldner et al., 2003).

In contrast, co-localization of GN-loop>J(3A) with ARF1 in BFA compartments was strongly reduced, thus resembling the strongly reduced accumulation of engineered BFA-resistant GNOM in ARF1-positive BFA compartments (Figure 6D). This result left unanswered the question of whether membrane association of GN-loop>J(3A) was reduced or the diminished BFA response was due to impaired ARF1 binding. Cell fractionation of seedlings revealed that GN-loop>J(3A) mutant protein partitioned between cytosol and membrane fraction like GNOM wild-type protein (Figure 6E and 6F). Membrane association of GNOM requires interaction of its DCB domain with the complementary fragment called ΔDCB, which is disrupted in the membrane-association-deficient mutant protein GNOM(B4049) (Figure 7) (Anders et al., 2008b).

**Figure 7.**
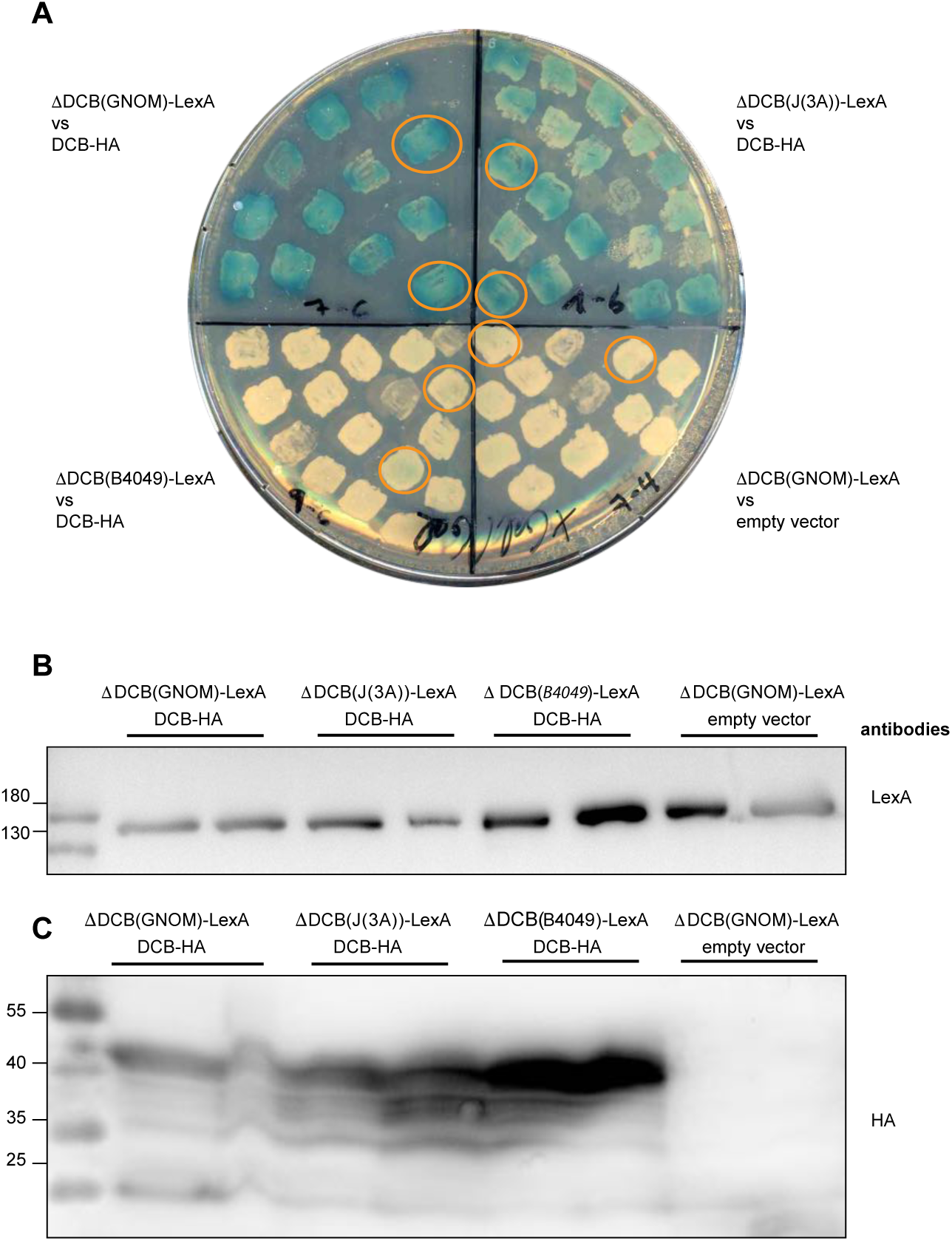
Y2H assay for DCB-ΔDCB interaction of GN-loop>J(3A). **(A)** ß-galactosidase activity stain. Unlike *gnom-B4049* (negative control, *lower left*), GN-loop>J(3A) (J(3A)) displayed DCB-ΔDCB interaction (*upper right*). *Upper left*: GNOM wild-type, (positive control; *lower right*: empty-vector control. See also Grebe et al., 2000 and Anders et al., 2008b. (**B, C**) Expression levels of constructs used for the interaction assay (protein extracts from circled colonies in (A) detected by immunoblots with specific antisera indicated on the right: (B) LexA (DNA-binding domain) fused to ΔDCB domains of GNOM wild-type (GNOM) and mutant (J(3A), GN-loop>J(3A); B4049, GNOM-B4049) proteins; (C) HA-tagged transactivation domain fused with DCB domain of GNOM (DCB-HA).

A yeast two-hybrid assay of DCB-ΔDCB interaction was positive for GN-loop>J(3A), like GNOM wild-type and in contrast to GNOM(B4049) (Figure 7). In conclusion, several lines of evidence suggest that GN-loop>J(3A) has normal membrane-association activity and that its BFA insensitivity is consistent with reduced ARF1 binding.

GNOM like other ARF-GEFs forms dimers (Grebe et al., 2000; Anders et al., 2008b). Co-immunoprecipitation with anti-Myc or anti-GFP beads revealed that Myc-tagged GN-loop>J(3A) mutant protein was able to dimerize with GNOM-GFP wild-type protein (Figure 8A). However, endogenous ARF1 was only detected in the precipitate of anti-GFP beads, which suggested that ARF-GEF dimers consisting of GNOM wild-type protein and GN-loop>J(3A) mutant protein have the same impaired interaction with ARF1 as GN-loop>J(3A) homodimers (Figure 8A and 8B). This puzzling result is consistent with the observation that GN-loop>J(3A) failed to rescue both *gnom-emb30* and *gnom-B4049* mutants, as described above (see Figure 3 and 4; Supplemental Figure 1) (Anders et al., 2008b). These results suggest that strong interaction of ARF1 with its exchange factor requires simultaneous binding of two ARF1•GDP molecules by the two SEC7 domains of the ARF-GEF dimer.

**Figure 8.**
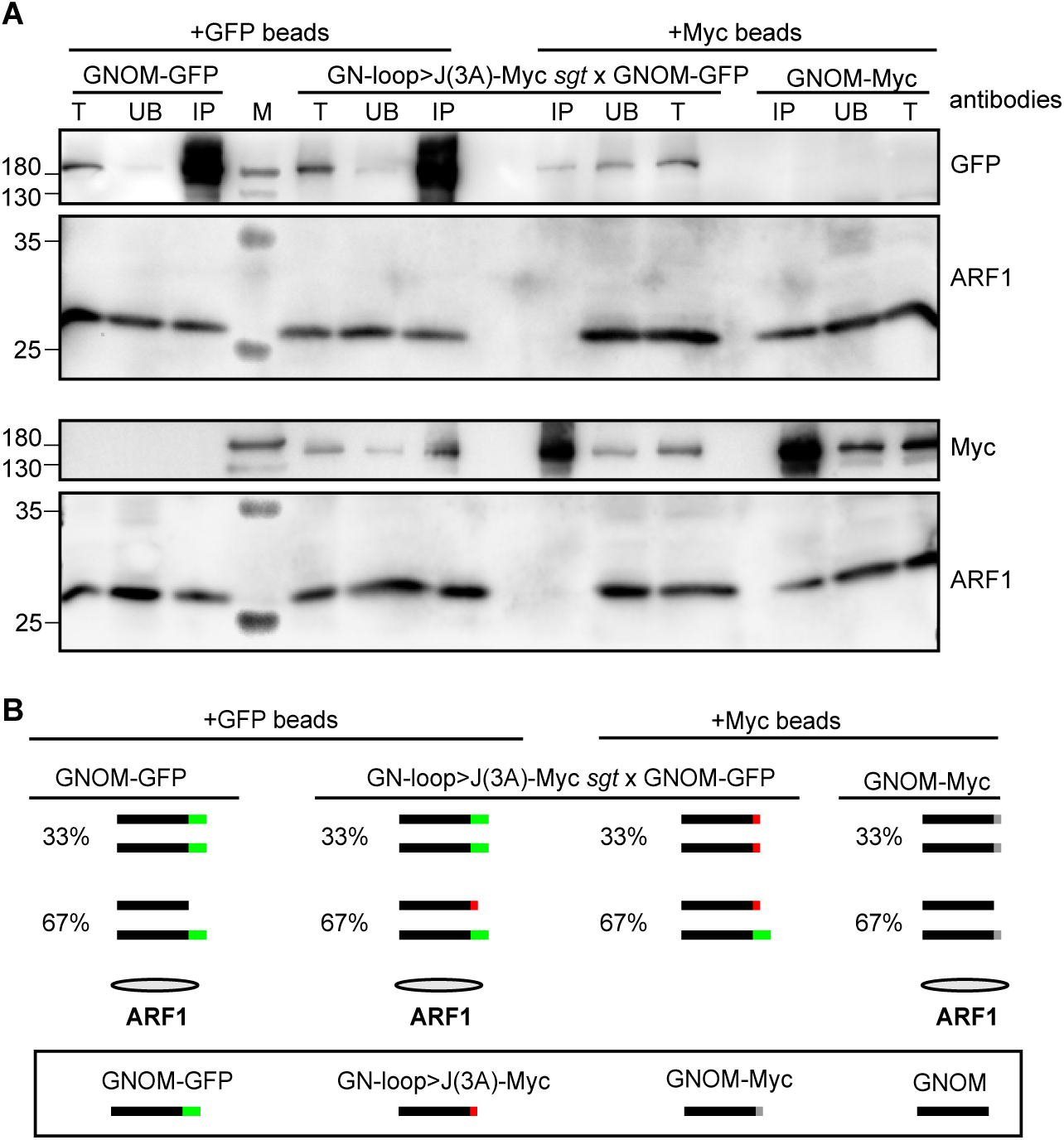
ARF1 binding by ARF-GEF dimers. **(A)** Co-immunoprecipitation of GNOM-GFP and GN-loop>J(3A)-Myc from seed-ling extracts with either anti-GFP or anti-Myc beads, revealing inter-action of GNOM wild-type with GN-loop>J(3A)-Myc mutant protein but no ARF1 binding by GNOM heterodimer. Precipitates were probed with anti-GFP, anti-ARF1 and anti-Myc antisera. T, total extract; UB, unbound; IP, immune-precipitate; M, molecular markers (sizes in kDa, *left*). **(B)** Diagram of expected co-immunoprecipitation results showing precipitated GNOM dimers and ARF1. Tags: green, GNOM-GFP (wild-type); red, GN-loop>J(3A)-Myc (mutant); grey, GNOM-Myc (wild-type); no tag, endogenous GNOM.

### A model of ARF1 activation-dimerization and hydrolysis-monomerization cycle

We propose the following model of how ARF1 dimers required for the scission of membrane vesicles are generated (Figure 9). Cytosolic GDP-bound ARF1 molecules are monomeric. They interact with membrane-associated ARF-GEF dimers, with the loop after helix J of the SEC7 domain playing a critical role in ARF1 binding. Productive complex formation requires cooperativity, i.e. simultaneous interaction of two ARF1•GDP molecules with the two SEC7 domains of ARF-GEF dimers. As a consequence, two adjacent ARF1 molecules undergo conformational change, resulting in GDP-GTP exchange, membrane insertion of the myristoylated N-terminus, and direct physical interaction of the two adjacent ARF1•GTP molecules. Following vesicle scission, GAP-assisted hydrolysis of GTP would alter the conformation of ARF1, disrupting the dimer and releasing monomeric ARF1•GDP into the cytosol (Figure 9B). Considering the conservation of the overall domain organization of large ARF-GEFs (Casanova, 2007; Bui et al., 2009), it is highly likely that cooperative ARF1 binding by ARF-GEF dimers as a mechanism of forming active ARF1 GTPase dimers on the donor membrane applies to eukaryotes in general.

**Figure 9.**
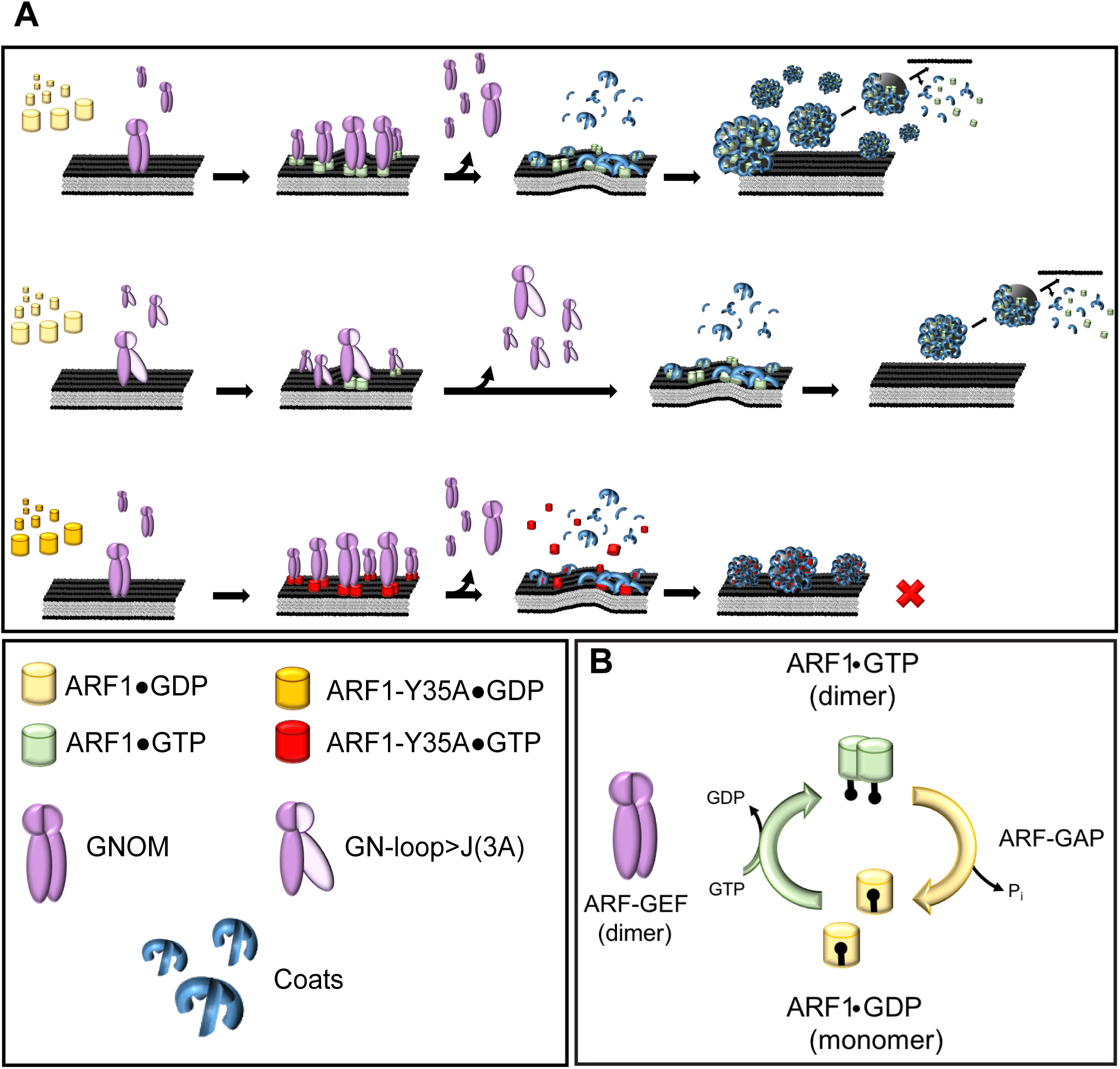
Model of ARF1 dimer formation. (A) Model of how ARF1 dimers required for scission of membrane vesicles are generated through simultaneous binding and activation by ARF-GEF dimers. ARF1 dimer formation by wild-type ARF-GEF dimers during GDP-GTP exchange on membrane (*top row*). GN-loop>J(3A) mutant protein reduces efficiency of ARF1 dimer formation because of reduced ARF1 binding (*middle row*). Two ARF1-Y_35_A proteins are each activated by ARF-GEF dimers but fail to interact, interfering with vesicle scission (*bottom row*). (B) Activation-hydrolysis cycle. Two ARF•1GDP monomers are simultaneously activated by membrane-associated ARF-GEF dimer, resulting in ARF1•GTP dimer. GTP hydrolysis facilitated by GTPase-activating protein (ARF-GAP) releases ARF1•GDP monomers from the membrane into the cytosol; P_i_, inorganic phosphate.

## Materials and Methods

### Plant material and growth conditions

Plants were grown under permanent light conditions (Osram L18W/840 cool white lamps) at 23°C and 40% humidity in growth chambers on soil or agar plates. Previously published lines that were used in this study: *gnom* mutant alleles *B4049*, *emb30-1, B4049/emb30-1* (Busch et al., 1996), *R5* (Geldner et al., 2004); transgenic lines expressing GNOM-Myc, GN-ML-Myc or GNOM-GFP from the *GNOM* promoter (Geldner et al., 2003), ARF1-YFP from *RPS5A* promoter and ARF1-T_31_N-YFP, ARF1-Q_71_L-YFP from an estradiol-inducible promoter (Singh and Richter et al., 2018). *gnom-sgt* mutant: The Ds-induced *sgt* allele was generated in an Ac-Ds mutagenesis experiment and isolated for its *gnom*-like mutant phenotype (insertion line SGT2467; Kumaran et al., 1999). The deletion on chromosome 1 eliminates nine genes from At1g13940 (5’ end of Ds) to At1g14020 (3’end of Ds) including *GNOM* (At1g13980; Figure 3B). For genotyping the following primers were used (Figure 3C and 3D): Primers for detection of *gnom-sgt* deletion (expected band size: 270 bp):

(N.A. 212) In At1g13940-sense: 5’ GGGGGGAGGGTATAAGAG 3’

(N.A. 213) DS-element-5’-antisense: 5’ ACGGTCGGGAAACTAGCTCTAC 3’

(N.A. 210) DS-element-3’-sense: 5’ GGTTCCCGTCCGATTTCGACT 3’

(N.A. 211) In At1g14020-antisense: 5’ AAGACACATGAGTGATTC 3’

Primers for detection of *gnom-sgt* homozygosity (expected band sizes: Myc-tagged *GNOM* transgene, 496 bp; endogenous *GNOM* gene, 373 bp; *gnom-sgt* homozygotes with transgene, 496 bp only):

(S.R.264) GNOM-over-tag-sense: 5’ GAAAGTGAAAGTAAGAGGC 3’

(S.R.263) GNOM-over-tag-antisense: 5’ CGTAGAGAGGTGTTACATAAG 3’.

Primers for detection of *B4049* mutation *in B4049/+* heterozygous seedlings (expected band sizes: allele

*B4049*, 535 bp; wild-type allele, no band):

(M.E.N.64) B4049-mut-for: 5’ TTAACAGGGATCCAAAGAATA 3’

(M.E.N.66) 3A-WT-rev: 5’ TCTGGAGTAGTCCTGATCTC 3’.

Primers for detection of wild-type *GNOM* in *B4049/+* heterozygous seedlings (expected band sizes: wild-type allele, 535 bp; allele *B4049*, no band):

(M.E.N.65) B4049-WT-for: 5’ TTAACAGGGATCCAAAGAATG 3’

(M.E.N.66) 3A-WT-rev: 5’ TCTGGAGTAGTCCTGATCTC 3’.

*emb30-1* genotyping:

A first PCR amplifying a 2496bp *GNOM* fragment including the *emb30-1* mutation site was performed with the primers:

(S.R.36) T391-S: 5’ TTCAAGTTCTCAATGAGTTTGCT 3’

(S.R.87) emb30-homo-AS: 5’ CTCACTTGTAAGGTCACGAACCAGTT 3’.

The first PCR product was used as a template for another PCR amplifying a 357bp *GNOM* fragment including the *emb30-1* mutation sites with the primers:

(S.R.36) T391-S: 5’ TTCAAGTTCTCAATGAGTTTGCT 3’

(S.R.37) T391-AS: 5’ CATTGTTGCAGATGGAGTGAA 3’.

The PCR product was cleaved using the restriction enzyme HinfI to distinguish the *emb30-1* mutant allele from the *GNOM* wild-type allele in the segregating *emb30/+* population. The expected restriction patterns are: wild-type allele, 193 bp, 65 bp, 47 bp, 32 bp, 20 bp; allele *emb30-1*, 193 bp, 97 bp, 47 bp, 20 bp.

#### Binary vector constructs, generation of transgenic plants and crosses

To generate the loop>J(3A) mutation, the amino acids residues 744, 745 and 747 (EI(R)T) were changed to alanines (AARA) by site-directed mutagenesis. Mutagenesis PCR was performed on the genomic fragment GNXbaI^wt^-Myc (Geldner et al., 2003) in pBlueScript, using the following primers:

Loop>J(3A) sense: 5’ AATGCGGCCAGGGCTACTCCAGAACAAGGTGC 3’

Loop>J(3A) rev: 5’ AGTAGCCCTGGCCGCATTGTTGCAGATGGAGTG 3’.

The *GN::GN-loop>J(3A)XbaI-Myc* fragment was cloned via XbaI into pGreenII(Bar) expression vector and transformed into *Arabidopsis thaliana* ecotype Col-0. T1 plants were selected using phosphinotricine. Four different transgenic lines showed good expression and two of them were chosen for further analysis.

*GN-loop>J(3A)-Myc* #5 was crossed into *sgt*, *B4049*, *emb30-1* backgrounds and analyzed for complementation. For co-immunoprecipitation analysis, *GN-loop>J(3A)-Myc* was crossed with

*RPS5A::ARF1-YFP* or *EST::ARF1-T_31_N-YFP*, and *GN-loop>J(3A)-Myc* (*sgt* heterozygous background) was crossed with *GNOM-GFP*.

To generate an estradiol-inducible ARF1-YFP variant, site-directed mutagenesis was performed on pEntry-ARF1-T_31_N-YFP (Singh and Richter et al., 2018), using the following primer combination:

ARFA1C-WT-MUT-S: 5’ [Phos]GCTGGTAAGAcgACTATCCTcTACAAGC 3’

ARFA1C-WT-MUT-AS: 5’ AGCATCGAGACCAACCATC 3’.

The Y_35_A mutation was introduced into pEntry-ARF1-YFP, pEntry-ARF1-TN-YFP and ARF1-QL-YFP by site-directed mutagenesis using the following primers:

ARFA1C-Y_35_A-MUT-S: 5’ [Phos]TACTATCCTCgcaAAGCTCAAACTTGGAGAGATC 3’

ARFA1C-Y_35_A-MUT-AS: 5’ TTCTTACCAGCAGCATCG 3’.

To generate RFP-tagged ARF1 variants, the CDS of RFP with a N-terminal AvrII restriction site was amplified and cloned into pDONR221 (Invitrogen) generating a pEntry clone. The RFP gene and part of the KAN resistance gene of the pEntry clone were then introduced via AvrII and SspI restrictions sites into the YFP-tagged ARFA1c Entry clones mentioned above, thereby replacing the YFP tag. The different ARF1 fragments were then introduced into a modified ß-estradiol-inducible pMDC7 vector by gateway LR reaction (Singh and Richter et al., 2018).

### Cloning of constructs for transient expression in protoplasts

CDS of ARFA1C, ARFA1C-T_31_N and ARFA1C-Y_35_A were amplified from pEntry clones mentioned above by using Sense-Primers containing NheI restriction site and Antisense-Primers containing BamHI restriction site. Amplified ARF fragments were introduced into pFK059 (Singh and Richter et al., 2018) via NheI and BamHI restriction sites.

NheI-ARFA1C-S: 5’ gatctcgctagcATGGGGTTGTCATTCGGAAAGTT 3’

BamHI-Stop-ARFA1C-AS: 5’ ggcagtggatccCTATGCCTTGCTTGCGATGTTGT 3’.

### Physiological tests

For primary root growth assays, 50 five-days old seedlings were transferred to agar plates containing 10 μM BFA for 24h and seedling growth was analyzed using ImageJ software. Gravitropic response of 50 five-days old seedlings was measured by ImageJ software after transferring seedlings to 10 μM BFA plates and rotating them by 135° for 24h. Lateral root primordia formation was analyzed after transferring 7-day old seedlings for 3 days on 20 μM NAA-containing agar plates and clearing the roots (Geldner et al., 2004). To examine the vasculature of 7 to 10-days old cotyledons, seedlings were shaken for several hours in 3:1 ethanol/acetic acid solution at room temperature (Geldner et al., 2004). Light microscopy images were taken with Zeiss Axiophot microscope, Axiocam and AxioVision_4 Software. Image size, brightness and contrast were edited with Adobe Photoshop CS 3 Software.

### Yeast two-hybrid interaction assays

Assay and constructs of GNOM-DCB (aa 1-246), GNOM-ΔDCB (aa 232-1451) and GNOM-ΔDCB(B4049) (aa 232-1451; G579R) were as described (Grebe et al., 2000; Anders et al., 2008b). GNOM-ΔDCB(J(3A)) was generated by site-directed mutagenesis using primers mentioned above.

### Quantitative transport assays

Protoplasts were prepared and electrotransfected as previously described (Künzl et al. 2016). Harvesting and analysis of medium and cell samples as well as calculation of the secretion index was performed as described (Bubeck et al., 2008). ARF1 expression levels were evidenced indirectly by detection of GFP. Both *ARF1* and *GFP* coding sequences are under control of the bidirectional *mas* promoter (consisting of a *mas1’* and a *mas2’* part) on the same plasmid, with *mas1’* directing GFP expression and *mas2’* directing ARF1 expression in a ratio of 1:10.

### Whole-mount immunofluorescence staining

Four to six-days old seedlings were incubated in 24-well cell-culture plates for 1 hour in 50 μM BFA (Invitrogen, Thermo Fisher Scientific) containing liquid growth medium (0.5x MS medium, 1% sucrose, pH 5.8) at 23°C and then fixed for 1 hour in 4% formaldehyde in MTSB at room temperature. Whole-mount immunofluorescence staining was performed manually as described (Lauber et al., 1997) or with an InsituPro machine (Intavis) (Müller et al., 2008). All antibodies were diluted in 1x PBS buffer. The following antisera were used for immunofluorescence staining: mouse anti-c-Myc mAb 9E10 (Santa Cruz Biotechnology) diluted 1:600; rabbit anti-ARF1 (Agrisera) diluted 1:1000; rabbit anti-AtγCOP (Agrisera) diluted 1:1000; anti-mouse Alexa488 (Invitrogen) and anti-rabbit CY3 (Dianova)-conjugated secondary antibodies were diluted 1:600. Nuclei were stained with DAPI (1:600 dilution).

### Confocal microscopy and processing of images

Fluorescence images were acquired with the confocal laser scanning microscope TCS-SP2 or SP8 from Leica, using a 63x water-immersion objective and Leica software; Zeiss LSM880 with Airy Scan and Zeiss software was also used. Overlays and contrast/brightness adjustments of images were performed with Adobe Photoshop CS3 software. Intensity line profiling was performed with the respective software.

### FRET-FLIM analysis

Four-to-five days old seedlings were incubated 4-6hours in liquid growth medium containing 20µM estradiol. FRET-FLIM measurements were performed at Leica TCS-SP8 upgraded with the rapidFLIM system from Picoquant (TimeHarp 260 time-correlated single-photon counting (TCSPC) module). YFP was excited with a pulsed 470 nm diode laser (LDHPC470B) with a 40 MHz pulse frequency. Emission was recorded at 495–550 nm by a HyD SMD detector until reaching a count of 1000 photons per pixel. Data were analyzed using the SymPhoTime software. n-exponential reconvolution with a mono-exponential decay function was used to fit the TCSPC histograms against a measured instrumental response function (IRF) (Fäßler and Pimpl, 2017; Mehlhorn et al., 2018). Box plots of measured fluorescence lifetimes were generated by using the box plot tyreslab web tool (http://boxplot.tyerslab.com; Spitzer et al., 2014). Statistical significance was calculated using a two-sample Student’s t test. Measurements were taken from at least 5 different seedlings in epidermal cells near the differentiation zone of the root.

### EM analysis

Four-to-five days old ARF1-YFP and ARF1-Y35A-YFP seedlings were incubated in liquid growth medium containing 10μM estradiol for 4h. For ultrastructural analysis, 1-1.5 mm long seedling root tips were high-pressure frozen, freeze-substituted in acetone containing 2.5% OsO_4_ and embedded in epoxy resin. Ultrathin sections were stained with uranyl acetate and lead citrate and viewed in a Jeol JEM-1400plus TEM at 120 kV accelerating voltage. For more information, see (Singh and Richter et al., 2018).

### Subcellular fractionation

2g of plant material were ground in liquid nitrogen and suspended in 1:1 extraction buffer (50mM Tris pH 7.5, 150mM NaCl, 1mM EDTA, 1mM PMSF) supplemented with protease inhibitors (cOmplete EDTA-free®, Roche). Of cell lysates, 100 µl were taken as total fraction (T). Then cell lysates were cleared by centrifugation at 10,000 × *g* for 10 min at 4°C and 100 µl of supernatant (S10) were saved for further analysis. The pellet was dissolved in 1 ml extraction buffer and 100 µl were frozen (P10). After 1x 100.000 × *g* centrifugation at 4°C for 1h, 100 µl supernatant (S100) were stored and the pellet was dissolved in 200 µl extraction buffer of which 100 µl were stored (P100*). 25μl of 5x Lämmli buffer were added to 100 μl samples.

### Co-immunoprecipitation analysis

The immunoprecipitation protocol was modified from ref. 31. 3-5g of 8 to 10-days old Arabidopsis seedlings were homogenized in 1:1 lysis buffer containing 1% Triton-X100. Seedlings bearing estradiol-inducible ARF1-YFP, ARF1-T_31_N-YFP or ARF1-Q_71_L-YFP were incubated in 20μM estradiol-containing liquid MS Media with sugar for 7h. For immunoprecipitation, anti-Myc-agarose beads (Sigma) or GFP-Trap beads (Chromotek) were incubated with the plant extracts at 4°C for 2h and 30 min. Beads were then washed twice with wash buffer containing 0.1% Triton-X100 and three times without Triton-X100. Bound proteins were eluted by boiling the beads in 2x Lämmli buffer at 95°C for 5min. Twice the usual amount of beads was used for immunoprecipitation involving GN-loop>J(3A)-Myc with anti-Myc beads or ARF1-YFP with anti-GFP beads.

### SDS-PAGE and protein gel blotting

SDS-PAGE and protein gel blotting with PVDF membranes (Millipore) were performed as described (Lauber et al. 1997). All antibodies were diluted in 5% milk/TBS-T solution. Antibodies and dilutions: mouse anti-c-Myc mAb 9E10 (Santa Cruz Biotechnology) 1:1000, mouse anti-GFP (Roche) 1:2500, rabbit anti-calnexin (Agrisera) 1:2000, rabbit anti-AALP (anti-aleurain (Holwerda et al., 1990)); a gift from Inhwan Hwang) 1:1000, rabbit anti-ARF1 (Agrisera) 1:2500, rabbit anti-SEC7 (Steinmann et al., 1999) 1:2500, mouse anti-LexA (Santa Cruz Biotechnology) 1:1000, POD-conjugated anti-HA (Roche) 1:4000, anti-mouse (Sigma) or anti-rabbit peroxidase-conjugated (Merck Millipore) or alkaline phosphatase-conjugated antibodies (Jackson Immuno Research) 1:10000. Detection was performed with the BM-chemiluminescence blotting substrate (Roche) and FusionFx7 imaging system (PeqLab). Image assembly was performed with Adobe Photoshop CS3, and ImageJ software was used for quantification of relative protein amounts.

## Acknowledgements

We thank Tobias Pazen and Rebecca Stahl for technical assistance, Inwhan Hwang for providing published material, and Martin Bayer, Jeff Dangl, Christopher Grefen, Niko Geldner and Thorsten Nürnberger for discussion and critical reading of the manuscript.

## Author contributions

Conceptualization: G.J.; Investigation: S.B., M.E.N., S.R., H.B., Y.-D.S., M.K.S., A.-M.F.; Formal analysis: S.B.; Resources: V.S.; Visualization: S.B.; S.R.; Project administration: S.B., G.J.; Writing – original draft: G.J.; Writing – review & editing: S.B., M.E.N., S.R., H.B., Y.-D.S., M.E.N., A.-M.F., V.S., G.J.; Funding acquisition: G.J., M.E.N., Y.-D.S.

## Data and materials availability

All data are available in the manuscript or the supplementary materials. Materials are available on request from G.J.

---

**Supplemental Figure 1.**
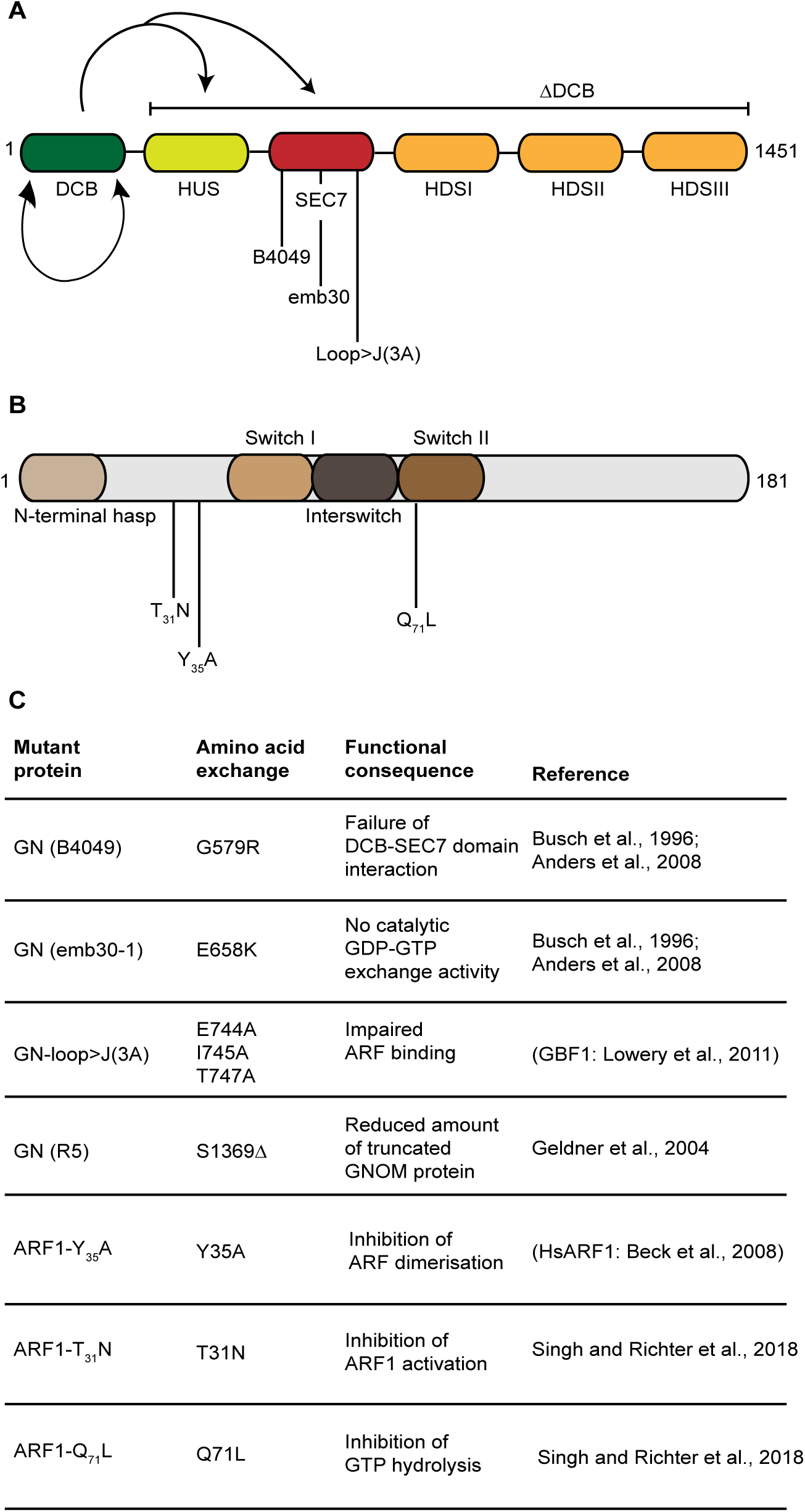
Overview of mutant variants of ARF-GEF GNOM and ARF1 used in this study. (**A, B**) Schematic presentations of domain architecture for GNOM (A) and ARF1 (B) with relevant mutations indicated. The DCB domain (green) interacts with another DCB domain and with both HUS domain (yellow) and catalytic SEC7 domain (red) of the complementary ΔDCB fragment (arrows). Not drawn to scale: GNOM, 1451 aa; ARF1, 181 aa. (**C**) Functional consequences of amino acid exchange in mutant proteins.

